# LiverZap: A chemoptogenetic tool for global and locally restricted hepatocyte ablation to study cellular behaviours in liver regeneration

**DOI:** 10.1101/2023.03.13.531762

**Authors:** Elizabeth M. G. Ambrosio, Charlotte S. L. Bailey, Iris A. Unterweger, Jens B. Christensen, Marcel P. Bruchez, Pia R. Lundegaard, Elke A. Ober

## Abstract

The liver restores its mass and architecture after injury. Yet, investigating morphogenetic cell behaviours and signals that repair tissue architecture at high spatiotemporal resolution remains challenging. We developed LiverZap, a tuneable chemoptogenetic liver injury model in zebrafish. LiverZap employs formation of a binary FAP-TAP photosensitiser followed by brief near-infrared illumination inducing hepatocyte-specific death and recapitulating mammalian liver injury types. The tool enables local hepatocyte ablation and extended *live* imaging of regenerative cell behaviours, critical for studying cellular interactions at the interface of healthy and damaged tissue. Applying LiverZap, we show that targeted hepatocyte ablation in a small region-of-interest is unexpectedly sufficient to trigger Liver Progenitor-like Cell (LPC)-mediated regeneration, challenging the current understanding of LPC activation. Associated dynamic bilary network rearrangement and E-cadherin relocalisation suggest modulation of cell adhesion as an integral step of LPC-mediated liver regeneration. This precisely targetable live cell ablation model will enable addressing of key regeneration paradigms.

## Introduction

Functional recovery after liver after injury relies on the capacity to re-establish organ mass and tissue architecture. Liver regeneration studies have focused mostly on the source of cell replenishment and the molecular pathways of mass (Forbes and Newsome, 2016; Michalopoulos, 2007; So et al., 2020). However, how the hepatic tissue architecture is restored, and the critical morphogenetic behaviours of the constituent cell types that mediate liver regeneration are still poorly understood.

Liver function depends on the intricate organisation of all hepatic cell types (Arias et al., 2020). Hepatocytes carrying out the main metabolic functions of the liver interact with both the vascular and biliary networks (Figure 1A). Biliary epithelial cells (BECs) compose the biliary network transporting hepatocyte-produced bile to the gallbladder and intestine. Upon mild or acute liver injury, such as partial hepatectomy, the remaining hepatocytes drive tissue mass restoration by hypertrophy and proliferation (Michalopoulos, 2007; Miyaoka et al., 2012). However, if hepatocyte proliferation is impaired, for instance in chronic liver injury, BECs can contribute new hepatocytes in mammals and zebrafish. In this process, BECs de-differentiate into Liver Progenitor-like Cells (LPCs), often referred to as oval cells in mammals, which can give rise to hepatocytes and BECs (Choi et al., 2014; He et al., 2014; Manco et al., 2019; Raven et al., 2017; So *et al*., 2020). The appearance of LPCs is associated with an expansion and remodelling of the biliary network (Choi *et al*., 2014; Kamimoto et al., 2016; Kamimoto et al., 2020; Kok et al., 2015). However, the significance of this remodelling and how it contributes to liver regeneration remains unclear. Elucidating the dynamic cellular processes of liver damage and repair requires both extended *live* imaging and high spatial resolution. Despite significant progress with intravital imaging (Vats et al., 2021) extended *live* imaging is still challenging in rodents (Cheng et al., 2021). In contrast, zebrafish are ideally suited for live imaging of cell and tissue dynamics, given their transparency and fast organ formation. The cellular composition, liver functions, and key cellular pathways controlling liver development and regeneration are largely conserved between zebrafish and mammals (Oderberg and Goessling, 2023; So *et al*., 2020; Wang et al., 2017). Yet, the majority of available zebrafish hepatocyte ablation models, such as the Nitroreductase-Metronidazole (NTR-MTZ) system, require prolonged treatments with pharmacological agents (e.g. 24-36 hours of Metronidazole; Curado et al., 2008). Moreover, MTZ toxicity can sensitise the larvae(Mathias et al., 2014) and render them incompatible with long-term *live* imaging. Furthermore, regeneration initiates concomitant with ongoing cell ablation in these models, which hinders the distinction of the two processes. Finally, investigating the interface between regenerating and healthy tissue, relevant to human disease in which tissue damage is heterogeneous across the liver(Martini et al., 2023) is not possible using global hepatocyte ablation systems. Optogenetic approaches (Varady and Distel, 2020) provide both precise spatial and temporal control of tissue manipulation. Genetically encoded photosensitisers, molecules releasing reactive oxygen species (ROS) upon illumination, can impair cell functions via photo bleaching-induced damage or cell death (Bulina et al., 2006). Yet *in vivo*, efficient ablation and activation in deeper tissues varies between different photosensitisers, representing the biggest challenges for optogenetic ablation models (Liu et al., 2021). The recently developed binary FAP-TAP system comprises a genetically encoded fluorogen-activating protein (FAP) that binds the malachite green derivate dye MG-2I to form a targeted and activated photosensitiser (TAP). Near infrared (NIR) illumination then initiates ROS production, that has been shown to induce cell death *in vitro* and in zebrafish cardiomyocytes and cause mitochondrial damage upon organelle-specific targeting in neurons (He et al., 2016; Missinato et al., 2021; Xie et al., 2020). The superior penetration properties of NIR light are ideal for effectively targeting deep tissues such as the liver, including reduced off-target excitation of endogenous chromophores (Deliolanis et al., 2008).

**Figure 1.**
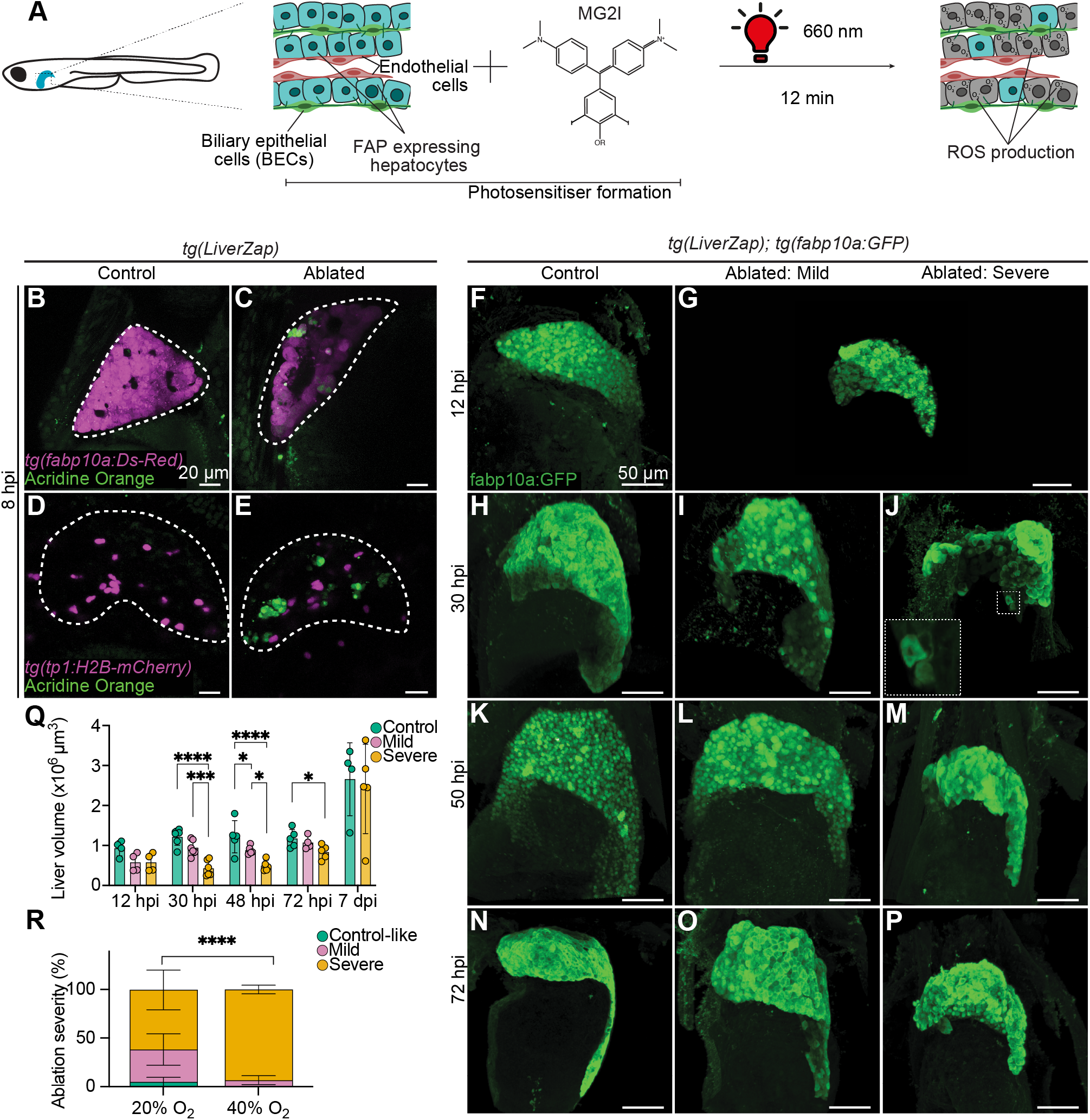
LiverZap activation elicits hepatocyte death and can be modulated. **A**. The LiverZap system expresses the photosensitising protein dL5** in hepatocytes. Upon combination with the dye MG-2I and NIR activation, ROS production triggers cell death. **B-E** Following NIR illumination, acrindine orange (green) marks apoptotic hepatocytes (B-C), while BECs remain unaffected (D-E); 5µm maximum intensity projections are shown. **F-P** Maximum intensity projections showing whole *tg(LiverZap);tg(fabp10a:eGFP)* livers at 12, 30, 50 and 72 hpi after LiverZap activation. Magnification in J shows detached cells. **Q** Quantification of liver volume of ablated LiverZap-and control livers. Statistical significance was determined by 2-way ANOVA followed by Tukey’s multiple comparison test (*p < 0.05, ***p < 0.001, ****p<0.0001); (N=2, n>3). **R** Distribution of ablation phenotype after LiverZap activation under normoxia or 40% O_2_ concentration. Statistical significance was determined by Chi-Square (****p<0.0001; N=3, n>30 per repeat).

Adapting FAP-TAP to the zebrafish liver, we developed LiverZap, a powerful chemoptogenetic tool for targetable hepatocyte ablation. Based on its optogenetic nature and a very short activation time, it is suitable for extended *live* imaging, and separating injury and repair processes. Spatially restricted LiverZap activation enables the study of local liver injury, relevant for elucidating human liver disease.

## Results

### LiverZap-mediated hepatocyte ablation can be modulated

To generate a chemoptogenetic hepatocyte ablation tool for extended *live* imaging, we developed LiverZap by adapting the FAP-TAP system (He *et al*., 2016). FAP expression, encoded by dL5**, was placed under the control of hepatocyte promoter *fatty acid-binding protein 10a (fabp10a)*. In combination with MG-2I dye FAP forms an active photosensitizer (FAP-TAP). Importantly, MG-2I was washed prior to illumination with NIR light at 660 nm generating ROS (Figure 1A). Stable transgenic *tg(fabp10a:dL5**-mCer3)* larvae, abbreviated *tg(LiverZap)*, were crossed to *tg(fabp10a:dsRed)* to visualise hepatocytes and exposed to 12 minutes NIR light. Cell death was evaluated *live* 8 hours post illumination (hpi) by the membrane-permeable nucleic acid-binding dye acridine orange. Green acridine orange fluorescence and apoptotic bodies were detected in DsRed-expressing hepatocytes of *tg(LiverZap)* larvae incubated with MG-2I, compared to control larvae displaying no signal (Figure 1B-C). Terminal deoxynucleotidyl transferase dUTP nick end labeling (TUNEL) further determined apoptosis as main cell death mechanism, consistent with previous reports(He *et al*., 2016) (Figure S1D-E). To test if ROS production impacts adjacent cells, acridine orange incorporation was assessed in the *tg(tp1:H2B-mCherry)* reporter marking BECs by expressing H2B-mCherry under the Notch responsive element Tp1(Ninov et al., 2012). H2B-mCherry-positive cells were devoid of acridine orange signal (Figure 1D-E), validating that LiverZap triggers hepatocyte-specific cell death, without a bystander effect in adjacent cells.

Having established the hepatocyte-ablation capacity of LiverZap, we examined ablation efficacy and the regeneration process using tissue mass recovery as read-out. Liver injury was triggered in 5 days postfertilisation (dpf) *tg(LiverZap)* larvae and monitored liver volume over time by *tg(fabp10a:eGFP)* hepatocyte expression (Figure 1F-Q). At 12 hpi, hepatocyte ablation was ongoing according to a 20-40% liver reduction compared to non-ablated livers (Figure 1F-G,Q). Between 24 and 30 hpi, we observed two categories of hepatocyte ablation (Figures 1H-J, S1A-C): (i) mild ablation, encompassing about ≥30% smaller livers devoid of apparent organ morphology or tissue integrity defects, and (ii) severe ablation marked by about ≥70% liver size reduction accompanied by a loss of organ morphology, and severely disrupted tissue organisation, including detaching hepatocytes (Figure 1J). At 50 hpi, a 14% size gain in livers with severe hepatocyte ablation, demonstrated ongoing regeneration. Mildly ablated livers exhibited no significant change (Figure 1K-M, Q). At 72 hpi, size recovery was also detectable in mildly ablated livers, while severely injured livers showed a drastic volume increase, doubling their size in 24 hours (Figure 1Q). By 7 days post illumination (dpi), severely ablated livers had recovered to the control liver volume (Figure 1Q). Quantifying total hepatic cell numbers mirrored and corroborated liver regeneration dynamics (Figure S1F).

A prerequisite for elucidating the cell behaviours mediating injury and repair is to determine the precise time point when regeneration begins. Employing light sheet microscopy to visualise the regeneration process after LiverZap ablation, we imaged *tg(LiverZap);tg(fabp10a:eGFP)* larvae from 24 until 55 hpi, (Figure S1I, Video S1). GFP-positive hepatocytes die during the first hours, until maximum ablation is reached between 24 and 30 hpi, with only a few round and detached hepatocytes remaining (Figure S1I). From this point onwards, regeneration was evident by the restoration of liver mass and organ morphology. This demonstrates that LiverZap, due to its optogenetic design, is compatible with extended *live* imaging, allowing the investigation of dynamic tissue morphogenesis after injury.

Since the FAP-TAP system kills cells by ROS production, we tested whether altering oxygen availability from 40% to 20% (normoxia) oxygen could be used to control hepatocyte ablation. Quantification at 24 hpi, when mild and severe categories are distinguishable also by stereomicroscope (Figure S1A-C), revealed that hepatocyte ablation in ∼40% oxygen-containing medium was highly efficient, with ∼90% severely ablated livers, in contrast to ∼60% severely and ∼40% mildly ablated livers under normoxic conditions (Figures 1R, S1G). In contrast, shorter illumination is unsuitable for modulating LiverZap ablation, since solely the number of control-like larvae increased, instead of those displaying mild injury (Figure S1H).

### Severe LiverZap-induced hepatocyte ablation leads to a Liver Progenitor-like Cell response

Across species, the extent of hepatic injury decides which regeneration program restores liver mass (Forbes and Newsome, 2016; So *et al*., 2020). Like mammals, mild or acute injury in larval and adult zebrafish livers is resolved by hepatocyte proliferation, whereas BEC-derived LPCs will give rise to new hepatocytes after extensive hepatocyte death or inability of hepatocytes to proliferate (Choi *et al*., 2014; He *et al*., 2014; Huang et al., 2014; Oderberg and Goessling, 2023). Given that LiverZap produces both mild and severe hepatocyte ablation, we examined which regenerative response is elicited by employing histone inheritance for short term lineage tracing(Blanpain and Simons, 2013) similar to previous studies (Choi *et al*., 2014; He *et al*., 2014). We generated *tg(LiverZap);tg(fabp10a:eGFP);tg(tp1:H2B-mcherry)* fish to visualize hepatocytes by eGFP expression, while Notch-active BECs express tp1:H2B-mCherry (referred to as mCherry; Figure 2A-A’’’). Since mCherry is fused to the histone protein H2B, progeny derived from a mCherry^high^ cell inherit half of the fluorescent protein. Hence, if a Notch-active BEC/LPC gives rise to a Notch-inactive hepatocyte, the latter will inherit half the mCherry and be identified as mCherry^low^ cell, while Notch-active cells will maintain mCherry^high^ expression. As regeneration progresses at 48 hpi, control and mildly ablated livers show mutually exclusive GFP and mCherry expression, marking hepatocytes and BECs, respectively (Figure 2A-B’’’). Notably, in severely ablated livers, there are two populations of mCherry expressing cells: mCherry^high^ and mCherry^low^ cells. The latter are all fabp10a:eGFP-positive, suggesting their LPC origin and hepatocyte identity (Figure 2C-C’’’). This indicates that regeneration of mildly ablated livers occurs by hepatocyte proliferation, whereas in severely ablated livers, BECs de-differentiate into LPCs that can give rise to new hepatocytes. To distinguish whether BECs/LPCs proliferate before they differentiate into new hepatocytes or after to replenish the BEC population, cell-type specific proliferation was determined by 5-Ethynyl-2’-deoxyuridine (EdU) incorporation in *tg(LiverZap);tg(fabp10a:eGFP);tg(tp1:H2B-mCherry)* over time. In severely ablated livers, surviving hepatocytes proliferate first, at 12 hpi (Figure 2D), followed by BEC proliferation which peaks at 30 hpi (Figure 2E), and lastly both LPC-derived and remaining hepatocytes proliferate at 48 hpi (Figure 2D, F). Proliferation of all cell populations returned to control levels by 72 hpi (Figure 2D-F). Quantification of mitotic cells by staining for Phospho-histone-3 (PH3) during regeneration in control and severely ablated samples showed similar dynamics (Figure S2B-D). These findings show that BEC/LPC proliferation precedes differentiation of new hepatocytes and that the LPC-response does not deplete the BEC pool at any time (Figure S2A). In mildly ablated livers, hepatocyte proliferation increased after ablation at 12 hpi and 30 hpi and decreased to control-levels by 72 hpi (Figure 2D). Interestingly, at 12 hpi and 30 hpi, BECs also showed mildly increased proliferation (Figure 2E) suggesting BECs respond although they do not progress into LPCs. Together, LiverZap can elicit different ablation severities and distinct repair responses, recapitulating the predominant regeneration modes in zebrafish and mammals.

**Figure 2.**
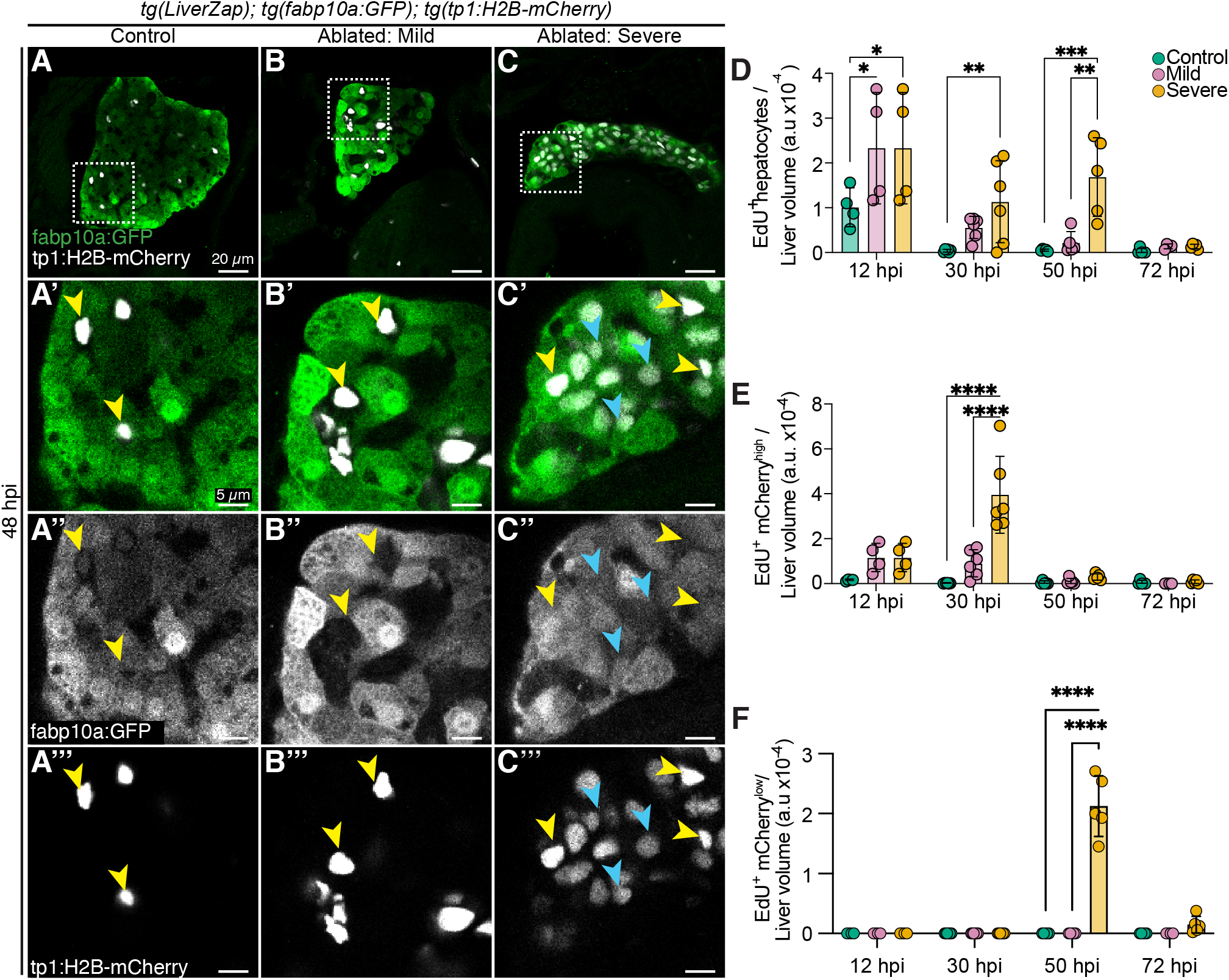
Severe liver injury after LiverZap activation triggers the LPC program. **A-C’’’** Expression of *tg(Tp1:H2B-mCherry)* marking BECs (grey) and *tg(fabp10a:eGFP)* highlighting hepatocytes (green) is mutually exclusive in control and mildly ablated livers (A-B’’). In severely ablated livers LPC-derived hepatocytes show low (cyan arrowheads) mCherry while BECs show high mCherry (yellow arrowheads) expression. Dotted squares in A, B and C indicate corresponding magnifications shown in A’-C’’’; 5µm maximum intensity projections. **D-F** Cell type-specific quantification of EdU incorporation in mild and severely LiverZap ablated livers (N=2, n>3). Statistical significance was determined by 2-way ANOVA followed by Tukey’s multiple comparison tests. (*p < 0.05, **p < 0.01, ***p < 0.001, ****p<0.0001).

### Spatially restricted LiverZap activation induces local LPC-mediated regeneration

The zebrafish NTR-MTZ liver regeneration models elicit organ-wide tissue damage, while locally restricted hepatocyte ablation cannot be recapitulated (Curado *et al*., 2008), which is a limitation considering that clinically relevant hepatic injury is often regionally restricted (Martini *et al*., 2023). We therefore tested whether restricted NIR illumination could activate LiverZap locally instead of globally. A three-dimensional region of interest (ROI) was illuminated at 660 nm in livers of *tg(fabp10a:DsRed);tg(LiverZap)* and *tg(tp1:H2B-mCherry);tg(LiverZap)* larvae using a confocal microscope with a tuneable white light laser (Figure 3A). At 17 hpi, acridine orange marking hepatocytes and loss of tissue integrity occurred exclusively in the illuminated ROI, representing about 10-20% of the liver volume (Figures 3B,C, S3A), whereas no acridine orange-positive BECs were detected (data not shown). Likewise, no unspecific hepatocyte death was detected in ROI-illuminated control livers (Figure 3B), showing that LiverZap can mediate spatially restricted hepatocyte ablation.

**Figure 3.**
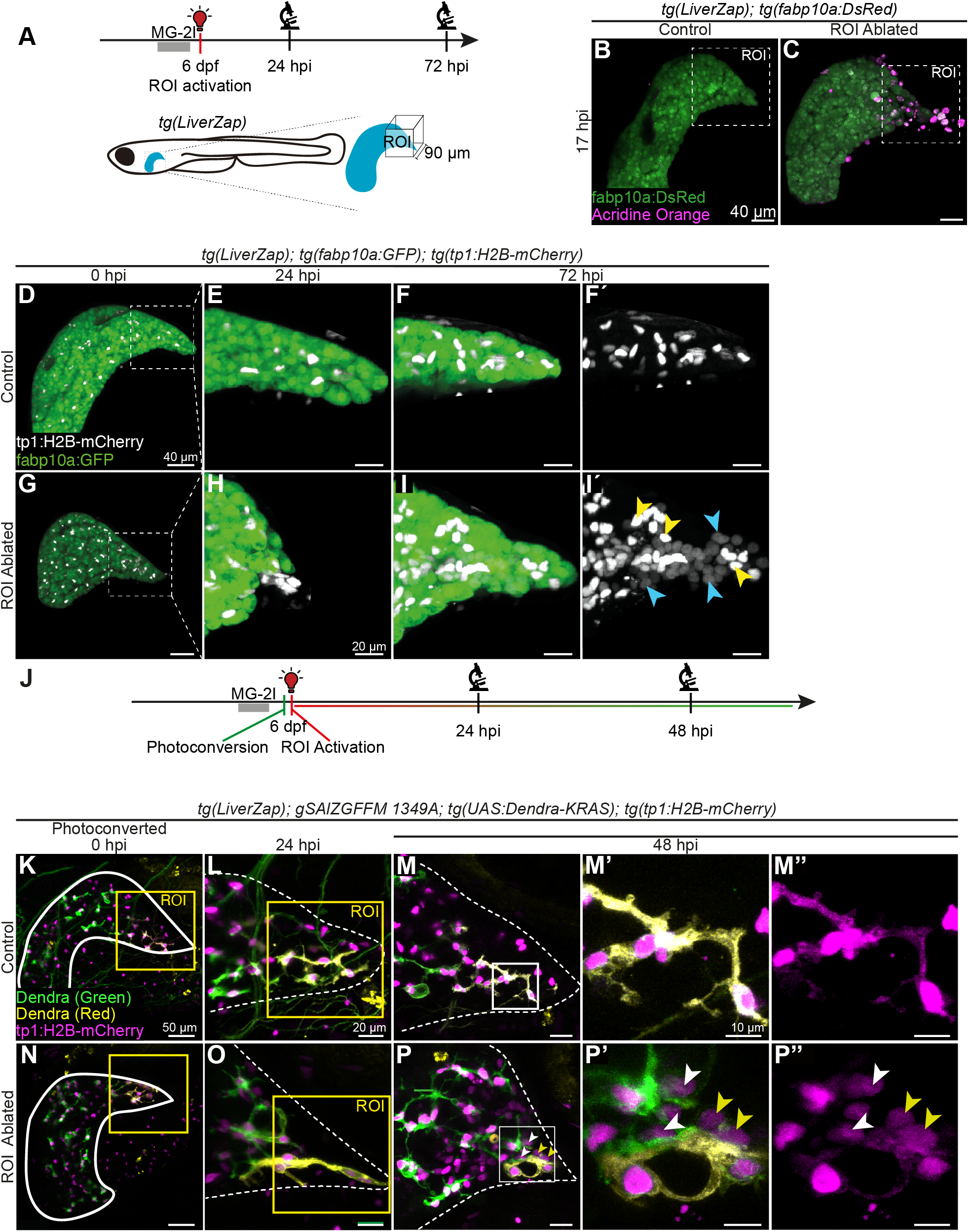
LiverZap ROI activation triggers a local LPC response. **A** Local activation of LiverZap using a confocal microscope WLL tuned at 660 nm. **B-C** Acridine orange (magenta) marks apoptotic cells after ROI LiverZap activation; hepatocytes express *tg(fabbp10a:eGFP)* (green). Maximum intensity projections show segmented livers. **D-I’** *tg(fabp10a:eGFP)-*positive hepatocytes (green) and *tg(tp1:H2B-mCherry)-*labelled BECs (igrey) were visualized after ROI LiverZap activation. At 72 hpi, LPC-derived hepatocytes are observed in ROI ablated livers expressing low mCherry (I’, blue arrowheads) while BECs have high mCherry (I’, yellow arrowheads). Masked maximum intensity projections are show. Dotted outline square in D and G depicts ROI shown in E-F’ an H-I’. **J** Dendra in BECs was photoconverted prior to LiverZap-mediated ROI ablation. Consecutive imaging monitored all specimen *live* at 24 and 72 hpi. **K-P** Dendra expression (green) in BECs was photoconverted prior to ROI LiverZap activation (yellow); BECs express *tg(tp1:H2B-mCherry)* in controls. (N=3,n>4). At 48 hpi, the ROI contains BEC-derived hepatocytes (mCherry^low^) showing converted (yellow arrowheads) or non-converted Dendra (white arrowhead). Maximum intensity projections are shown in K-L and N-O. 5µm projections are shown in M-M’’, P-P’’.

Addressing whether restricted hepatocyte ablation is sufficient to trigger regeneration, ROI-illumination was performed in *tg(LiverZap); tg(fabp10a:eGFP); tg(tp1:H2B-mCherry)* embryos (Figure 3D-I’
s). Similar to whole organ ablation, the source of new hepatocytes can be deduced from the inheritance of BEC-derived mCherry expression. Monitoring the ablation efficacy at 24 hpi by *live* imaging showed efficient hepatocyte ablation was associated with aggregation of the BECs in the ROI (Figure 3E,H), consistent with global ablation. By 72 hpi, hepatocytes had repopulated the ablated ROI (Figure 3I), indicating regeneration had been triggered. Remarkably, most hepatocytes in the ablated region, identified by fabp10a:eGFP and a characteristic round nuclear shape (Russell et al., 2019), exhibited mCherry^low^ expression (Figure 3I’), suggesting their BEC origin, and the activation of the LPC program. Imaging distant regions in the same ROI-ablated specimen revealed no apparent morphological changes, corroborating that the LPC program was exclusively activated in the illuminated ROI (Figure S3A-D’). This finding is very surprising, since current understanding in the field links the generation of new hepatocytes from BECs/LPCs with global hepatocyte death.

### BECs outside the ablated region contribute to the local LPC response

ROI-restricted LiverZap activation elicited the LPC response locally in a comparative small volume and appears to recover tissue mass within 72 hrs. This raised the question whether only BECs within the illuminated ROI gave rise to new hepatocytes or other cell sources also contributed. To exclusively trace BECs in the ROI, we turned to a photoconversion strategy (Arrenberg et al., 2009). To specifically express photoconvertible Dendra in BECs, we identified the gene trap line gSAlzGFFM1349A expressing the activator Gal4FF (Asakawa and Kawakami, 2009) in BECs (EMAG, KK and EAO unpublished) and combined it with the effector line UAS:Dendra-KRAS. Photoconverted Dendra was specific to BECs and still detectable after 72 hours (Figure S3E-J). In *tg(LiverZap);tg(tp1:H2B-mcherry); gSALzGFFM1349A; tg(UAS:Dendra-KRAS)* larvae, photoconversion was immediately followed by LiverZap activation in the same ROI (Figure 3J). Photoconversion and NIR light illumination of control larvae without MG-2I caused no injury (Figure 3K-M’’). Tissue morphology alteration concomitant with hepatocyte death and the condensation of BEC nuclei was apparent 24 hpi after ROI LiverZap activation (Figure 3O). At 48 hpi, BEC/LPC-derived hepatocytes appeared, distinguishable by their mCherry^low^ expression and hepatocyte-typical round nuclear shape (Figure 3P-P’’). Some of the LPC-derived hepatocytes had inherited converted red Dendra, highlighting new hepatocytes derived from BECs/LPCs in the ablated ROI (Figure 3P-P’’). Strikingly, a subset of BEC/LPC-derived-hepatocytes (mCherry^low^) showed solely non-converted green Dendra (Figure 2P’-P’), indicating that in addition to BECs/LPCs in the ablated region, BECs within an average distance of 88 µm in adjacent uninjured tissue (Figure S3L) respond to the injury, de-differentiate into LPCs and contribute to replenishing new hepatocytes after local injury.

### E-Cadherin decrease at BEC membranes is linked to injury induced biliary network collapse

The cellular behaviours and morphological changes elicited by liver injury and driving regeneration are largely unknown. Given the crucial role of BECs in regeneration, understanding the morpho-dynamic processes related to the biliary network, such as the de-differentiation of BECs into LPCs is of particular interest.

First, to quantify the change of biliary network structure during injury and regeneration, distances to the nearest neighbour were measured in livers using BEC nuclei following LiverZap activation in 5 dpf *tg(LiverZap);tg(tp1:H2B-mCherry)* larvae (Figure 4A-J). In control livers, BEC nuclei (mCherry^high^) are at all analysed time points distributed throughout the organ, with 40% having the closest neighbour at a 0-8 µm distance and 50% at 8-16 µm, respectively. In contrast at 12 hpi, midway through ablation, BEC nuclei commence clustering with 60% displaying a 0-8µm distance. This distribution is maintained in mildly ablated livers at 30 hpi, the maximum ablation timepoint. Severely ablated samples, however, show more extensive clustering of BEC nuclei as 85% are 0-8 µm apart. This doubling of the small distances compared to control, demonstrates increased clustering of BECs during injury. At the network scale, biliary Anxa4 expression at 24 hpi corroborates these morphological changes (Figure S4A-D). As regeneration is initiated, biliary network organization is progressively restored. By 72 hpi, mildly ablated livers show a BEC inter-nuclear distance distribution that is comparable to control larvae. Severely ablated livers at 72 hpi show increased distances between BECs, since 40% show a surprisingly larger distance than 24 µm between neighbours, which is rare in control livers, suggesting network restoration is still ongoing (Figure 4J). These data show that gradual condensation of BECs accompanies progressive hepatocyte death and provides a quantitative baseline for future functional studies of BEC network remodelling during injury and regeneration.

**Figure 4.**
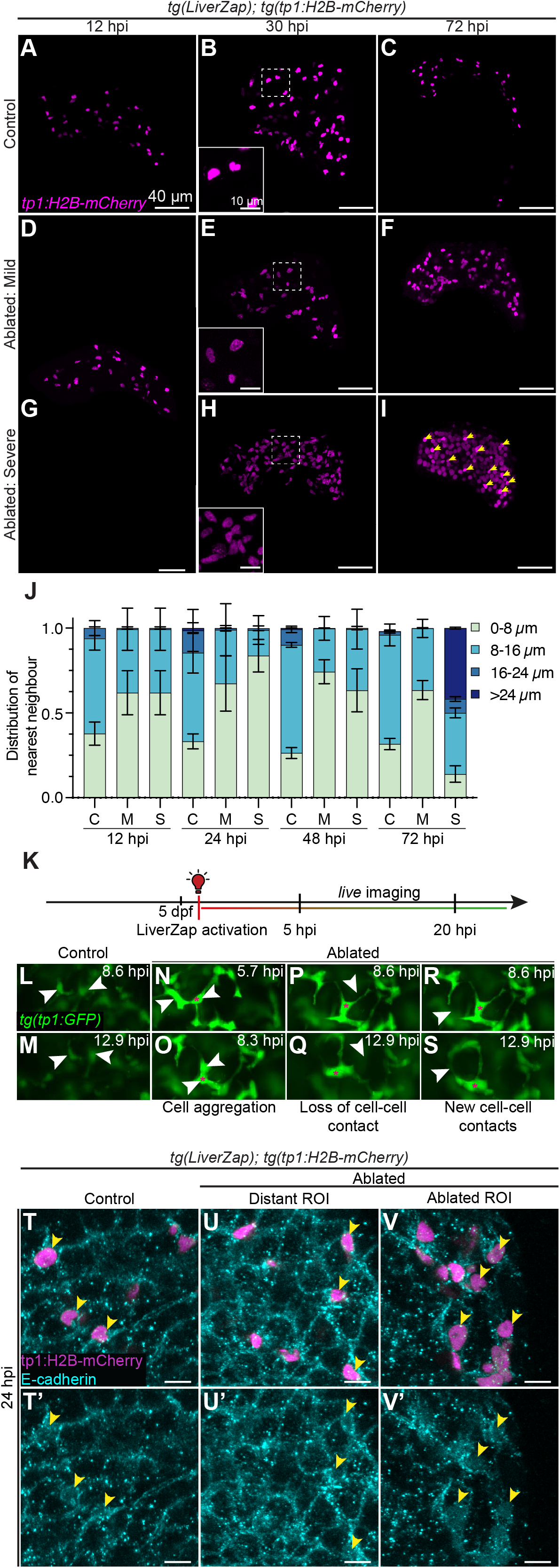
Injury-induced biliary network aggregation includes dynamic cell and E-Cadherin relocalisation. **A-I** *tg(tp1:H2B-mCherry)*-positive nuclei (magenta) serve as proxy for biliary network remodelling during ablation and regeneration. 10 µm masked maximum intensity projections are shown. **J** Distribution of the nearest neighbour distances displayed by mCherry^high^ nuclei populations in controls and following LiverZap ablation (N=2, n>3); C=Control, M=Mild, S=Severe. **K** Schematic showing sequence for live imaging experiment. **L-S** Selected frames from time-lapse imaging (VideoS2) of *tg(tp1:eGFP)-*expressing BECs (green) in control (L-M) and LiverZap-ablated livers (N-S). BECs exhibit dynamic cellular behaviours including cell aggregation (N-O), loss (P-Q) and gain of new cell-cell contacts (R-S) during the phase hepatocytes die. Magenta asterisks mark the same cell across the entire timelapse. **T-V’** E-cadherin expression (cyan) in control livers (T-T’), distant ROI (U-U’) and ablated ROI (V-V’). *tg(tp1:H2B-mCherry)* visualizes BECs (magenta). 10 µm maximum intensity projections shown. Yellow arrowheads highlight selected BECs.

The dynamic nature of BEC aggregation and re-distribution raised the question whether this is the result of active remodelling or whether the loss of the scaffold normally provided by hepatocytes could lead to a passive collapse of the network. *Live* imaging of *tg(tp1:eGFP)* in LiverZap-ablated livers during the injury phase from 5 to 20 hpi, revealed highly dynamic BEC behaviours, including cell rearrangements and filopodia-like extensions, while control BECs appear mostly static (Figure 4K-S; Video S2).

The finding that BECs become more dynamic as the biliary network remodels raised the possibility that cell adhesion needs to be adjusted to enable the dynamic behaviours facilitating network remodelling as BECs adopt an LPC state. Cadherins are candidate adhesion factors, and in particular E-Cadherin is highly expressed in mammalian BECs and required for maintaining biliary network integrity in mice (Nakagawa et al., 2014; Sasaki et al., 2001). Like in mammals, E-cadherin is expressed in BECs and at a lower level in hepatocytes, clearly outlining their cellular shapes in control livers and unablated ROIs (Figure 4T-U’). Upon ROI-injury, E-Cadherin was lost from hepatocyte membranes, in line with cell death induced integrity loss and hepatocyte detachment. In BECs, E-cadherin changed from the cell membrane to a largely intracellular localisation (Figure 4V-V’). Similar relocalisation occurred following global LiverZap activation (Figure S4 E-H’), suggesting reduction of cell adhesion as a general mechanism contributing to the aggregation of the biliary network after hepatocyte loss.

## Discussion

Here, we introduce LiverZap, a new chemoptogenetic hepatocyte ablation tool for studying liver injury in zebrafish. LiverZap ablation triggers severity-dependent regeneration programs corresponding to those in zebrafish and mouse models (Forbes and Newsome, 2016; Michalopoulos, 2007). LiverZap, does not require long pro-drug incubation and has a short activation time. It thereby complements and adds to the existing liver regeneration toolbox by (i) its compatibility with extended *live* imaging to investigate dynamic restorative cell behaviours, (ii) the possibility to temporally separate ablation and regeneration phases, and (iii) the capacity for eliciting regionally restricted hepatocyte cell death, enabling *live* studies of the interface between healthy and damaged tissue. Using LiverZap, we show that the biliary network undergoes dynamic remodelling accompanied by E-Cadherin relocalisation and cellular adhesion changes, suggesting a key role of network condensation in the de-differentiation process from BECs to LPCs.

### Chemoptogenetically encoded LiverZap can activate different regenerative responses

LiverZap can elicit both mild and severe hepatocyte ablation which reflect the two main modes of liver regeneration depending on the type and extent of injury. In zebrafish, it has been shown that following mild injury in larvae (Choi *et al*., 2014) or partial hepatectomy in adults (Kan et al., 2009; Sadler et al., 2007), the regenerative potential relies on the proliferation of differentiated hepatocytes. In contrast, upon extensive injury or impaired hepatocyte proliferation, BECs can de-differentiate to give rise to the transient LPCs. These have the potential to contribute to both hepatocytes and BECs (So *et al*., 2020). Short-term lineage tracing of BECs progeny and proliferation studies revealed that solely hepatocyte proliferation restores liver mass after mild LiverZap ablation, while severe LiverZap ablation triggered BEC de-dfferentation into LPCs, giving rise to new hepatocytes. This process is accompanied by distinct, cell-type specific proliferation phases: hepatocytes proliferate first, then BECs/LPCs and finally, LPC-derived hepatocytes. Both the cellular sources as well as the proliferation dynamics resemble those of other injury models including the widely used NTR/MTZ system in zebrafish (Choi *et al*., 2014; He *et al*., 2014). LiverZap findings will therefore be comparable and extend the currently available experimental tools.

A key advantage of LiverZap is that incubation with the MG-2I dye to form the FAP-TAP photosensitiser occurs prior to the short 12-minute ROS-inducing NIR illumination. This contrasts with the long drug treatments concomitant with cell ablation in the existing NTR-MTZ models (Choi *et al*., 2014; He *et al*., 2014; Huang *et al*., 2014), prone to obscuring regeneration onset. The short activation allows for the clear separation of injury and regeneration phases, evident by the drastic size reduction in the first 30 hpi, and subsequent recovery. Moreover, the LiverZap ablation mechanism allows circumventing possible unspecific MTZ effects on larval survival (Mathias *et al*., 2014) that can be detrimental to extended *live* imaging. This is clearly shown by *live* imaging for ≥30 hours and allowed to determine dynamic BEC behaviours and cell-cell interactions essential for de- and reconstruction of the liver architecture following injury.

### ROI activation of LiverZap elicits a local LPC response

Under physiological conditions, liver damage is often locally restricted, for example non-alcoholic fatty liver diseases originate pericentrally in human, while autoimmune hepatitis starts periportally (Martini *et al*., 2023). Regenerative processes consequently take place at the junction of healthy and diseased tissue. With LiverZap precise and restricted ROI activation allows to investigate these processes *live* and at high resolution. Extraordinarily, ROI hepatocyte ablation of 10-20% liver volume activated the LPC program in a regionally restricted fashion, suggesting that LPC activation does not exclusively happen after extensive, global hepatocyte death as previously shown (Choi *et al*., 2014; He *et al*., 2014; Manco *et al*., 2019; Raven *et al*., 2017; So *et al*., 2020). Instead, it suggests that the ratio of lost hepatocytes to BECs in a defined area may be sufficient to elicit this repair program. This could also explain recent observations of small foci of LPC-derived hepatocytes in NTR-MTZ mediated injury in adult zebrafish (Oderberg and Goessling, 2023). Therefore, we propose that LPC activation may be a consequence of BECs sensing their environment and reacting to massive hepatocyte cell death in their immediate surroundings. Elucidating the cellular morphological changes as well as the cell-cell signalling occurring at the interface of injured and healthy tissue will be crucial to compare with behaviours in hepatic disease such as cirrhotic nodules, where proliferation is restricted by fibrotic tissue and is therefore insufficient to restore liver function (Forbes and Newsome, 2016).

Furthermore, we show that restricted hepatocyte ablation activates the LPC program in BECs beyond those located in the ablated region, as BECs up to 100 µm (∼3-5 hepatocyte diameters) into the unablated tissue also contribute new hepatocytes. This strongly suggests that BECs sense and respond to signals from their environment, potentially through sensing molecules released by dying hepatocytes (Brenner et al., 2013; Eguchi et al., 2014; Jung et al., 2010). In addition, network remodelling associated with a LPC-response could also transduce mechanical cues beyond the actual injury border.

### Biliary remodelling accompanies hepatocyte cell death

Nuclear clustering and biliary network remodelling occur after extensive LiverZap and NTR-MTZ-triggered hepatocyte death (Choi *et al*., 2014), which is also a hallmark of mammalian ductular reaction (Kamimoto *et al*., 2016; Kaneko et al., 2015). Ductular reaction often precedes oval cell activation and is characterised by proliferation and expansion of the biliary compartment (Kamimoto *et al*., 2016; Kamimoto *et al*., 2020). While nuclear aggregation is more evident in severe samples, we show it also occurs in mild LiverZap-ablated livers, suggesting it may be the first response from BECs upon hepatocyte death, possibly due to local changes in the mechanical support. Therefore, extensive BEC network tissue remodelling, such as enhanced condensation of BECs, dominating in severely ablated livers, may serve as an early step in the LPC response. Similarly, ductular reaction in murine models can occur without the generation of new hepatocytes (Kamimoto *et al*., 2020) suggesting specific thresholds control the activation of the LPC program. Relocalisation of adherens junction component E-Cadherin from BEC membranes, implicates reduced cell adhesion and strongly supports this notion since mis-regulation of E-Cadherin levels is a hallmark of biliary diseases. BEC-specific deletion of E-Cadherin in mice causes cholangitis and cancer(Nakagawa *et al*., 2014), while in biliary atresia and intrahepatic cholangiocarcinoma it is aberrantly expressed in BECs and hepatocytes (Guedj et al., 2016). The loss of E-cadherin at BEC membranes is also reminiscent of epithelial-to-mesenchymal transition, where E-cadherin reduction and upregulation of N-cadherin results in acquisition of mesenchymal migratory behaviours (Lamouille et al., 2014). This, together with the observation of increased BEC protrusive activity further suggests that active cell rearrangement plays a key role in the remodelling of the biliary network upon liver injury In summary, LiverZap is a chemoptogenetic hepatocyte ablation model based on 12-minute NIR-light activation that enables (i) extended *live* imaging possibilities for capturing tissue remodelling during ablation and repair (ii) separation of injury and regeneration processes, and (iii) ROI hepatocyte ablation. LiverZap exhibits common hepatocyte- and LPC-mediated regenerative responses. Its experimental properties extend the existing regeneration toolbox and will enable analysis of dynamic cellular behaviours key for understanding how liver architecture is deconstructed during injury and rebuilt during repair.

## Acknowledgments

We thank all members of the Ober group for fruitful discussions, T. Piotrowski for invaluable support, K. Kawakami for the gSAlZGFFM1349A:Gal4FF line, D.Y.R. Stainier for the tg(UAS:Dendra-KRAS) line and S.E. Fraser for suggestions on oxygen concentrations. We further thank A. Vanoosthuyse for expert technical support, J. Bulkescher, A. Sheshtra and J. Dreier (DanStem Imaging core facility) for assistance with image acquisition, analysis and quantification, and the department of experimental medicine for expert fish care. We are grateful to Drs. A. Boni and P. Strnad (Viventis) for light sheet microscopy.

## Material and methods

### Zebrafish stocks

Adult zebrafish and embryos (*Danio rerio*) were kept and raised according to standard laboratory conditions (Westerfield, 2000). To prevent pigmentation, embryos were grown in E3 medium (5 mM NaCl, 0.17 mM KCl, 0.33 mM CaCl2, and 0.33 mM MgSO4) with 0.2 mM N-Phenylthiourea (PTU, Sigma-Aldirch cat # P7629), from 24 hpf onwards and under standard laboratory conditions. All experiments were performed according to ethical guidelines approved by the Danish Animal Experiments Inspectorate (Dyreforsøgstilsynet).

The following strains were used: *tg(EPV*.*Tp1-Mmu*.*Hbb:hist2h2l-mCherry)*^*s939*^ abbreviated *tg(tp1:H2B-mCherry)* (Ninov *et al*., 2012), *tg(EPV*.*tp1-Mmu*.*Hbb:eGFP)*^*um14*^ abbreviated *tg(tp1:eGFP)* (Parsons et al., 2009), *tg(−2*.*8fabp10a:eGFP)*^*as3*^ abbreviated *tg(fabp10a:eGFP)* (Her et al., 2003), *tg(UAS:Dendra-kras)*^*s1998t*^ abbreviated *tg(UAS:Dendra)* (Arrenberg *et al*., 2009), *tg(fabp10a:dsRed)*^*gz15*^ abbreviated *tg(fabp10a:dsRed)* (Dong et al., 2007) and gSAlZGFFM1349A:Gal4FF (Kawakami et al., 2010).

### Generation of transgenic lines

The fluorogen-activating protein dL5** (FAP) fused to monomeric cyan fluorescent protein mCer3 was excised from the pCS2+ MBIC5-mCer3 construct (He *et al*., 2016) and placed under the control of the *fabp10a* promoter (Her *et al*., 2003) by insertion into the *pHD157 - 2*.*8fabp10a:Cre;cryaa:Venus* construct (Ni et al., 2012). Stable transgenic carriers were generated by injecting 25-50pg DNA with I-SceI meganuclease into one-cell stage embryos as previously described (Soroldoni et al., 2009). Multiple transgenic founders were tested for hepatocyte ablation efficiency, of which two transgenic lines of similar efficiency were established. *Tg(fabp10a:dL5**-mCer3)*^*cph10 and cph11*,^ abbreviated *tg(LiverZap)* and are currently maintained in F6 generations. All experiments were carried out with heterozygous *tg(fabp10a:dL5**-mCer3)*^*cph10*^ larvae.

### Fluorescent immunostaining and imaging

Specifically, larvae were fixed in 4% paraformaldehyde (PFA) (Sigma Aldrich, cat# 6148) overnight at 4ºC. Embryos were then washed 3 × 5 minutes in PBST 0.1 % (0.1% Triton X-100 in PBS). The yolk was manually removed followed by permeabilisation with PBST 2% (2% Triton X-100 in PBS) for 1 hr at room temperature. Blocking and whole-mount antibody staining was performed in PBST 1% (1% Triton X-100 in PBS) with 10% serum (goat or donkey) and 1% Dimethyl Sulfate (DMSO, Sigma-Aldrich, Cat# D8418). Antibody incubation was done at 4ºC for 2 nights while gently shaking. Antibodies used included: mouse α-2F11 (1:1000; gift from Julian Lewis), goat α-Hnf4a (1:100; Santa Cruz, cat# sc-6556, lot# B1605), mouse α-E-cadherin (1:500; BD Biosciences, Cat# 610181, lot 9301695), rabbit α-pH3 (1:500; Millipore, Cat# 631257, lot 2825969), chicken α-GFP (1:500; Abcam, ab13970), rat α-mCherry (1:1000; Invitrogen, m11217 lot TL276838 and UJ287711). Fluorophore-conjugated secondary antibodies (Jackson Immunoresearch) were added at 1:500 to PBST 1% with 10% serum and 1% DMSO. To visualize nuclei, DAPI (Sigma-Aldrich, Cat# D9542) was added to the secondary antibody solution. Whole-mount immunostained embryos were mounted either in 1:2 Benzyl Alcohol:Benzyl Benzoate (BABB, Sigma-Aldrich cat# B6630 and 305197) clearing mixture or Vectashield (VectorLabs, Cat#H-1900) prior to imaging. The embryos were imaged either with a Leica SP8 or Leica Stellaris confocal microscope. Long term *live* imaging was performed using a LS1 Live light sheet microscopy system (Viventis Microscopy Sàrl). Images were processed with Imaris (Bitplane) image analysis software, Fiji(Schindelin et al., 2012) and Adobe Suite CS 2019.

### LiverZap activation

Embryos were incubated in E3 supplemented with 0.2 mM PTU and 500 nM MG-2I (synthesized as in He *et al*., 2016) protected from light, overnight or for a minimum of three hours at 28ºC, 20 or 40% O_2_. The next day larvae were transferred into a 6 cm diameter petri dish with fresh 20 or 40% oxygen enriched E3/PTU. The free-swimming larvae were placed under a near-infrared LED light at a 2 cm distance, inside a custom-built box (as described in He *et al*., 2016) equipped with an exhaustion fan to dissipate heat. Larvae were then illuminated for 12 minutes, and subsequently returned to the incubator at 28ºC with ambient oxygen levels. Control larvae without MG-2I underwent the same treatment. Ablation severity phenotypes were scored at 24 hours post illumination (hpi) under a fluorescent stereomicroscope based on liver size as described in results.

### Liver volume and nuclei segmentation

Imaris software was used to calculate the liver volume through manual segmentation of the liver using the “surface” module. Nuclear counts were determined with the “spots” module, including manual correction.

### Nearest neighbour quantification

Using the software Imaris (version preceding 9.0), all BECs were marked using the “spots” function. For each spot, the xyz coordinates were exported and employed to calculate the nearest neighbour distance using a custom Matlab script. Since the release of V9.0 of Imaris, this quantification can also be performed automatically through the software statistics tab.

### Live cell death analysis

Larvae were incubated in acridine orange (Sigma Aldrich, cat# A6014) (5µg/mL) in E3/PTU for 30 minutes protected from light at 28ºC and washed twice. Zebrafish larvae were anesthetised with 0.4% Tricaine in Tris (Sigma-Aldrich, Cat# A5040) and mounted in 1% low melting agarose (Lonza, Cat# 50080) supplemented with 0.16% tricaine. They were mounted on their left side to increase tissue accessibility and imaged using a Leica Stellaris confocal microscope.

### Cell proliferation assay

Embryos were incubated for 1hr at 28ºC with 400 µM 5-ethynyl-2’-deoxyuridine (EdU, Invitrogen Cat# A5040) and 5% DMSO in E3/PTU. The fish were then fixed with 4% PFA overnight at 4ºC with gentle shaking. EdU incorporation was visualised by using the Click-iT™ EdU Alexa Fluor™ 647 Imaging Kit (Invitrogen, Cat# C10340) prior to secondary antibody incubation.

### Region of interest LiverZap activation

Fish with the desired transgenic expression were incubated at 5 dpf with MG-2I as described for LiverZap activation. The next day, individual fish were transferred into fresh oxygen-rich E3/PTU (incubated overnight at 40% O_2_). The fish were anesthetised with 0.4% Tricaine and mounted on their left side in low melting agarose 1% in E3/PTU media supplemented with 0.16% tricaine. Once the agarose solidified, the chamber was filled with oxygen-rich E3/PTU with 0.16% Tricaine to prevent drying and maintain the anaesthesia. For ROI-ablation, fish were illuminated using a confocal microscope Leica Stellaris equipped with a White Light Laser (WLL) tuned at 660nm. Fish were imaged using a 40X water objective. The region of interest (ROI) was selected by digital zoom into the tissue (3X zoom). LiverZap activation in the ROI was achieved by using a laser speed of 10 Hz in combination with a low resolution of 256 × 256 pixels. The total Z-stack of the ROI was kept at 90µm illuminated at 5 µm intervals, which in the majority of cases encompasses the entire thickness of the tip of the left liver lobe. The WLL power was set up at either 85% or 100% and laser intensity at 100%. Each ROI was illuminated for a total of 20 or 40 minutes, equivalent to 5 or 10 cycles of imaging the complete z-stack. We observed an increasing need for laser power or imaging length as the laser aged from daily wear.

After illumination, individual larvae were carefully recovered from the agarose, placed in E3/PTU and ambient oxygen levels and kept in an incubator until the desired timepoint.

### Photoconversion and *live* imaging

Larvae were prepared and mounted in the same fashion as for ROI-LiverZap activation. For photoconversion, we illuminated the fish for 5 minutes with 10% laser power using a 405 nm laser in the Leica Stellaris confocal microscope. We simultaneously acquired both non-converted and converted Dendra signal and confirmed that all Dendra expressing cells were converted. To prevent unspecific photoconversion, imaging of non-converted Dendra was performed using very low 488 nm laser power (0.1%). We used a Leica Stellaris confocal microscope and acquired the images using a 40X water objective.

For *live* imaging with the LS1 Live light sheet microscope (Viventis), fish were mounted ventrally to improve tissue illumination and imaging acquisition. A 25X objective was used and acquisition was performed every 20 minutes.

### Experimental design and statistical analysis

Biological replicates represent different cohorts of larvae, bred on different days and/or different pairs and are indicated in each figure legend as N. Technical replicates are indicated with the letter n and represent individual embryos. Statistical analysis was performed using GraphPad Prism software and specific tests used are stated in figure legends.

